# Real-time label-free exploration of the dynamics and interactions of bacteriophages

**DOI:** 10.1101/2024.04.30.591896

**Authors:** Francesco Giorgi, Judith M. Curran, Liberty Duignan, Joanne L. Fothergill, Sam Chenery, Eann A. Patterson

**Affiliations:** Department of Materials Design and Manufacturing Engineering, University of Liverpool, The Quadrangle, Liverpool, L69 3BX, UK; Department of Clinical Infection, Microbiology and Immunology, University of Liverpool, Brownlow Hill, Liverpool, L69 3BX, UK, United Kingdom; Department of Mechanical and Aerospace Engineering, University of Liverpool, The Quadrangle, Liverpool, L69 3BX, UK

**Keywords:** bacteriophages, microscopy, bacteria, optical caustics, phage therapy

## Abstract

Bacteriophages are the focus of extensive research and monitoring their dynamics and interactions with bacterial hosts is crucial to characterise the mechanisms of infection and support potential applications in biotechnology and medicine. Traditional monitoring techniques rely on the fluorescent labelling of bacteriophages due to their size being nanometric. In this paper, we propose a novel, label-free method to generate optical signatures of bacteriophages in a conventional microscopy setup by exploiting the optical phenomenon of caustics. We utilised previously isolated *Pseudomonas aeruginosa* phage (pelp20 and phiKZ) and a novel *Escherichia coli phage* (EcoLiv25) to demonstrate detection and tracking within liquid laboratory media. The results obtained confirm the feasibility of visualising and monitoring over time a diverse array of bacteriophages, offering a simpler and less invasive means of observation for research and application in microbiology and related fields.

## 2. Introduction

Bacteriophages, commonly referred to as phages, are the most abundant form of microorganisms on Earth. These entities selectively infect and replicate within bacteria and play a pivotal role in regulating bacterial populations across diverse ecosystems, including soil, oceanic waters, and even the human body [1]. The morphology of phages is varied, with structures ranging from simple icosahedral heads attached to tails, which may be short and non-contractile, long and contractile, or even tailless. This diversity not only influences their size, typically within 20 nm to 300 nm, but also reflects their genomic diversity, with DNA contents varying significantly among different phage species [2]. This variability and the wide range of bacterial hosts they can infect underscore bacteriophages as subjects of study in microbiology, ecology, and biotechnology. Their potential applications include phage therapy as an alternative to antibiotics [3,4], as tools in molecular biology for genetic engineering [5], and as biocontrol agents in agriculture and food safety [6,7].

Given their nanometric size, the morphology of bacteriophages is typically characterized using high-resolution microscopy techniques such as Scanning Electron Microscopy (SEM), Transmission Electron Microscopy (TEM), or Atomic Force Microscopy (AFM) [8]. While powerful, these methods typically require complex and invasive sample preparation that can alter or destroy biological specimens like phages. Moreover, they do not allow for the real-time observation of phage dynamics or interaction with host cells in complex biological media, which is critical for understanding their behaviour, lifecycle, and interactions with bacterial hosts. In response to this challenge, the technique of single-virus tracking (SVT), also known as single-virus tracing, has been developed. SVT allows researchers to follow individual viruses at different parts of their life cycle, thereby providing dynamic insights into fundamental processes of viruses and interaction with host cells [9]. To enable SVT, strategies have been developed that involve staining phages with various fluorescent agents, from fluorescent proteins to quantum dots, which selectively highlight phages for dynamic monitoring over time [9-11]. Despite recent advances, fluorescence-based techniques introduce complications and uncertainty in results, as the staining process can affect the phage’s physical and biological properties, potentially altering their infectivity and behaviour. Fluorescent dyes and proteins can photobleach under intense light exposure, limiting observation times and complicating long-term studies of phage dynamics. Moreover, fluorescence microscopy requires the use of specific wavelengths of light, which can inadvertently damage biological samples or induce unwanted photochemical reactions [12]. These factors highlight the need for novel visualization techniques that facilitate non-invasive, real-time monitoring of bacteriophages within complex biological media, enhancing the accuracy of understanding their dynamics and interactions to facilitate the application of phages in real-life biotechnological and medical scenarios.

In this paper, we investigate the ability of bacteriophages to generate optical signatures many times their own size in an optical microscope by exploiting the optical phenomenon of caustics, thus enabling the real-time monitoring of their dynamics without the need for fluorescent labelling. Caustics, characterized by focused light patterns formed through the refraction or reflection of light, have found applications across various fields of science and engineering. Traditionally employed in engineering to track the formation and propagation of cracks in materials, caustics offers a non-invasive method to assess structural integrity [13]. Further applications include enhancing optical systems and guiding light in photonic devices [14,15]. The usage of caustics has been also extended to the nanoscale to visualize and track nanoparticles as small as a few nanometers in diameter without labelling or invasive preparation techniques [16]. Moreover, caustics-based optical microscopy has been used successfully in characterizing the dynamics of bacterial cells and the early stages of biofilm formation, demonstrating its potential as a tool for real-time and label-free monitoring of microorganisms [17].

## 3. Materials and Method

The study investigates three bacteriophage strains: EcoLiv 25, PELP20, and PhiKZ. EcoLiv 25 is a novel *E. coli* phage isolated by L. Duignan able to infect *E. coli* bacteria. The EcoLiv 25 solution was prepared by inoculating 10 ml of the host strain E. coli K-12 in Luria-Bertani broth (LB; Sigma Aldrich, US) at the mid-exponential stage of growth (OD600 of 0.5) with 100 µl of a pure phage stock. This mixture was incubated at 37°C for 12 hours and subsequently filtered to remove the host bacteria. The other phages, PELP20 and PhiKZ are known to target *P. aeruginosa* bacteria and were sourced from previously published studies [18, 19]. Phage titers were determined using plaque assay, with counts in the range of 10^14 to 10^15 plaque-forming units per milliliter (PFU/mL).

Phages were imaged in a standard optical inverted microscope (Axio Observer.Z1 m, Carl Zeiss), mounted on antivibration feet (VIBe, Newport, UK) to isolate the sample from the environment, using 60 μl of phage solution in a deep cavity (250 ± 10 μm in depth) in microscopy slides. The microscope was equipped with a monochrome camera (AxioCam ICm1, Carl Zeiss) to acquire images and record videos at up to 30 fps and with a stage-top incubation system (Incubator PM S1, Heating Insert P S1, Carl Zeiss) to maintain constant the temperature of the phage solutions at 37°C during the experiments. The resolving power of the optical setup was increased to generate caustic signatures of the phages by making some simple adjustments to the normal setup of the microscope following the procedure described by Patterson and Whelan [16], including setting up Kohler illumination, closing the aperture of the condenser lens to its minimum of about 1 mm diameter, and inserting a narrowband filter (546 nm) in front of the 100W Halogen lamp. To validate the label-free caustic signatures associated with phage in solution, fluorescence images of the phages were acquired with the same optical setup using a fluorescent light source (pE-300^white^, CoolLED, UK). Phages were stained with the fluorescent dye SYTO9 (Thermo Fisher Scientific, Waltham, US), a well-known green-fluorescent nucleic acid stain that can permeate the membranes of both eukaryotic and prokaryotic cells. Dyes of the SYTO family have already been used to successfully stain bacteriophages [20]. One microliter of SYTO9 solution was added to 100 μL of concentrated EcoLiv25 bacteriophage preparations. Bacteriophage solution was allowed to stain and equilibrate for 20 min at room temperature in the dark before being transferred into the microscope slide cavity for imaging. Single-phage tracking was performed by acquiring videos of the phages and evaluating their dynamics over time using the ImageJ plugin Trackmate, which provided a way to automatically segment spots or roughly spherical objects from a 2D or 3D image and track them over time [21].

## 4. Results and Discussion

### 4.1 Phages imaging and tracking

The morphologies of the phages were first characterized by acquiring TEM images of the four phage strains investigated in this study (Fig. 1). The images acquired show that all of the phages investigated are part of the *myoviridae* family, possess a long-contractile tail, and have an overall size ranging from approximately 200 nm for the Pelp20 phages to approximately 300 nm for the PhikZ phages. Following the morphological characterization, the imaging capability of the optical setup used in this study to visualize phages was validated against fluorescence microscopy by capturing consecutive images of the same region of the sample by switching from caustic mode to fluorescence mode, which involved changing the light source. Figure 2 shows that all of the phages identifiable in the images captured in the fluorescence mode are also present in the images captured in the caustic mode, confirming the ability of the caustic technique to generate optical signatures for phages and confirming the ability to perform real-time monitoring and characterization of their dynamics without any requirement for fluorescent labelling. Caustics of the three phage strains are shown in figure 3 and illustrate that they all produce similar optical signatures on the orders of microns (as previously reported in studies on nanoparticles [16]), proving the possibility to monitor phages of different size and genome. Given the cability of the optical setup to generate visible caustics from the phages tested, we proceeded to monitor and characterise their dynamics over time. As illustrated in Figure 4, phages in solution exhibit the random motion typical of brownian particles, caused by the collisions with the surrounding fluid molecules.

**Figure 1:**
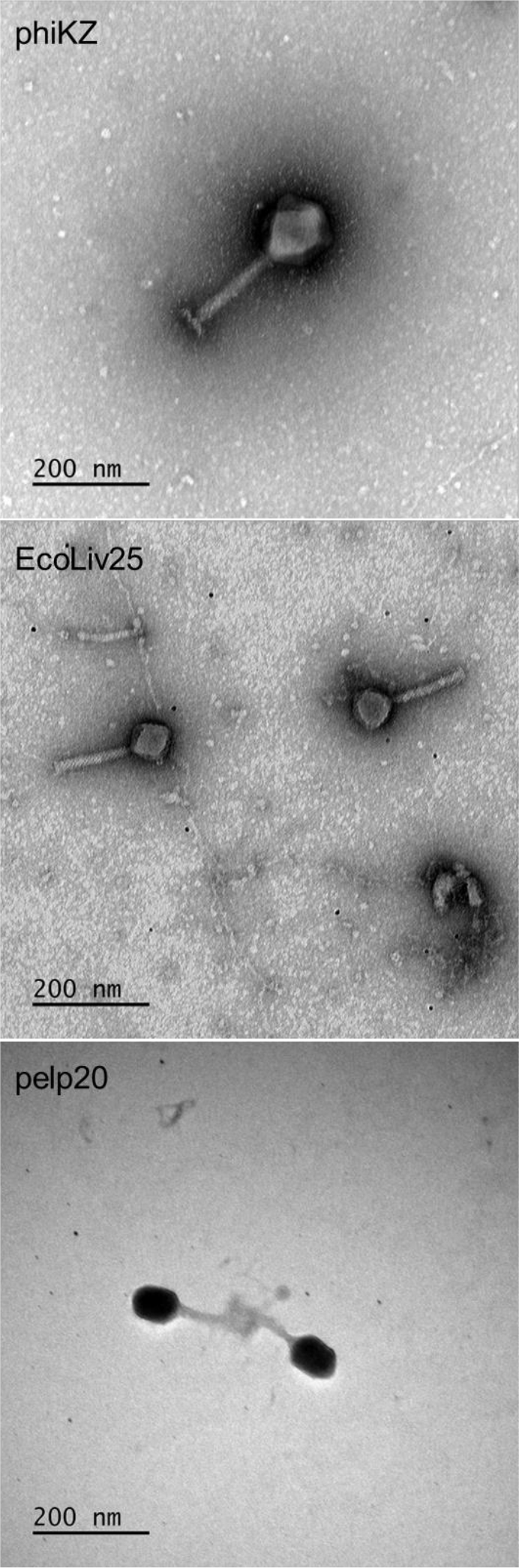
TEM images of the phage strains used in this study.

**Figure 2:**
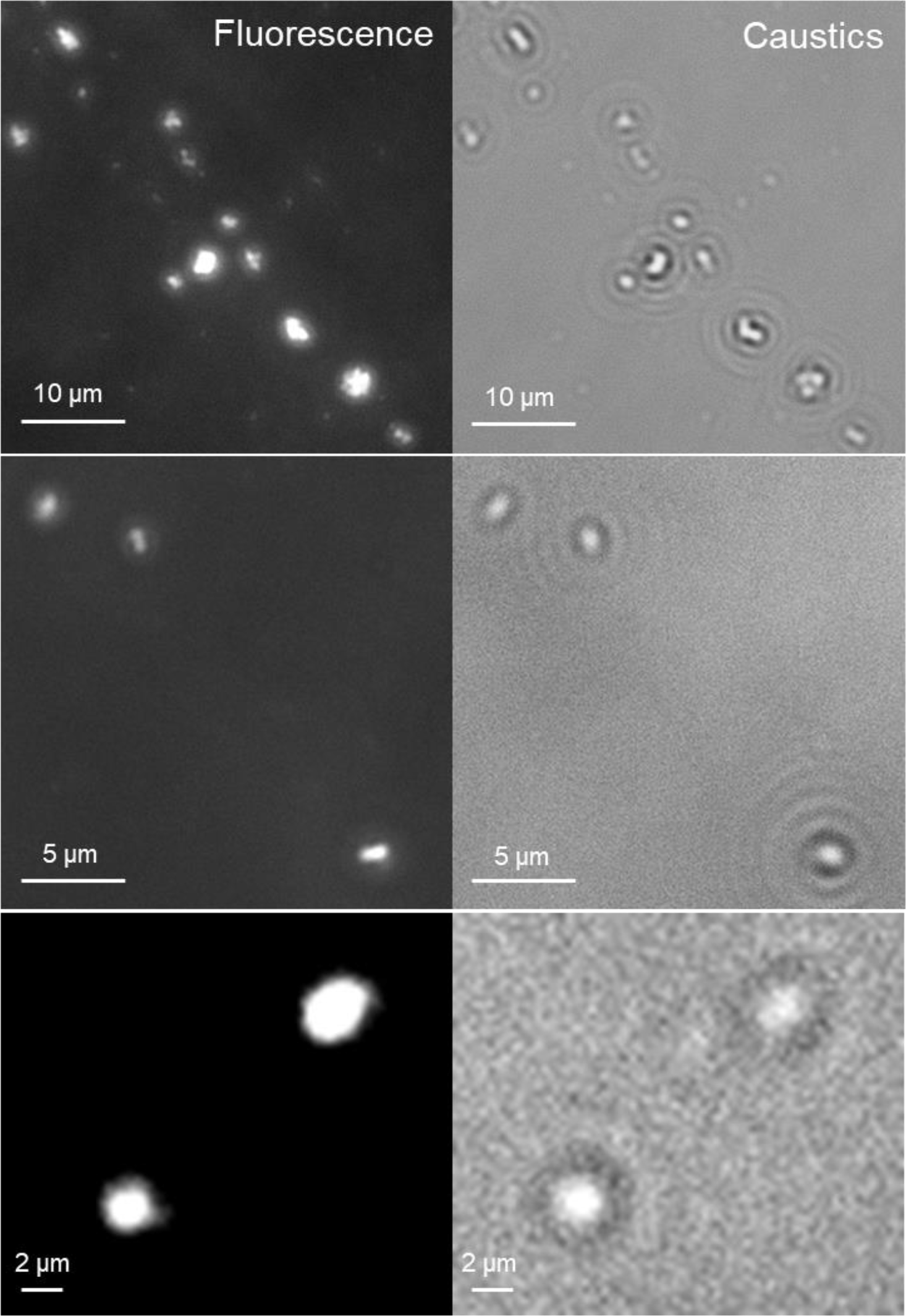
Comparison between the same EcoLiv25 phage populations imaged with an inverted optical microscope in fluorescence mode (left) and caustics mode (right). Images refer to three distinct regions of the sample and were acquired with different magnification from low magnification (top) to high magnification (bottom).

**Figure 3:**
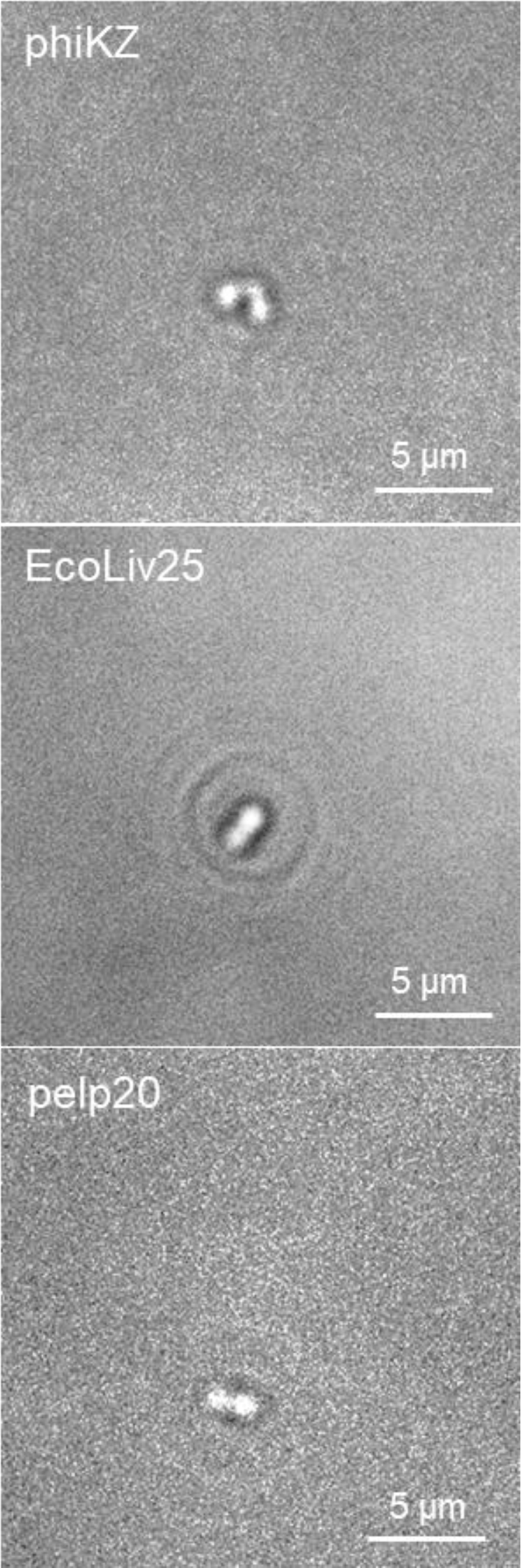
Caustics produced by label-free phages tested in this study. The optical signatures produced are bigger than the real phages in solution, allowing for their monitoring without the need of fluorescence labelling.

**Figure 4:**
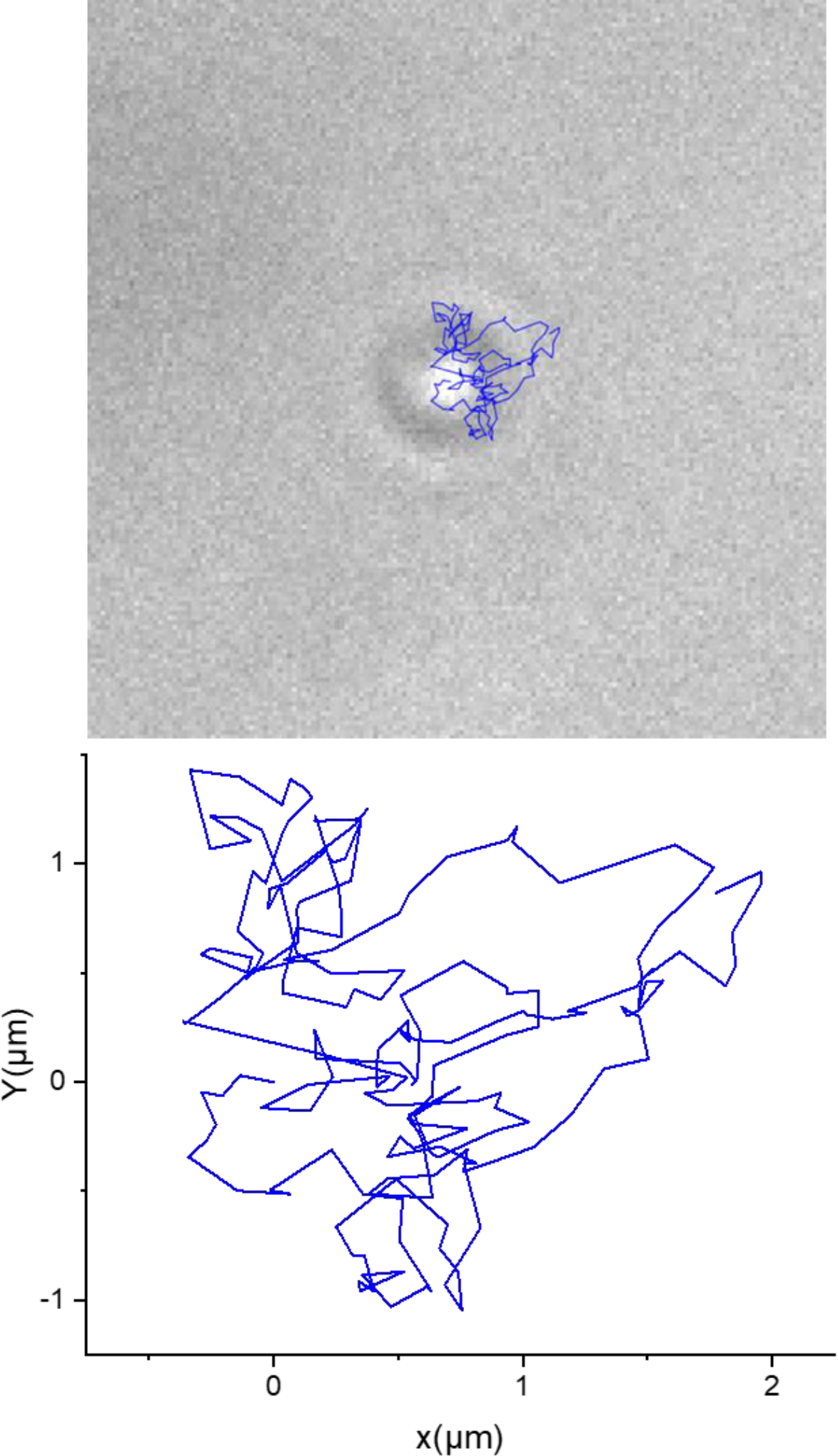
2D random dynamics (blue line) of a pelp20 bacteriophage monitored for a period of 5 seconds (top) and a plot of the same dynamics (bottom).

### 4.2 Phages – bacteria interaction

Building on our ability to visualize bacteriophages and to characterize their motion in real time using caustics, we extended our investigation to understand the potential to capture the interactions between bacteriophages and their bacterial hosts. Preliminary results, illustrated in Figure 5, provide an early indication of the possibility to observe these interactions and the mechanism of infection by analyzing the optical signatures of both the bacteria and the interacting bacteriophages. The comparison of the optical signatures between non-exposed bacteria and those exposed to bacteriophages reveals observable differences, which could be tentatively attributed to the attachment of bacteriophages to the external membrane (Fig. 5b) and/or to the compromised membrane integrity of the infected bacteria (Fig. 5c). However, while these differences suggest a promising avenue both for distinguishing between infected and non-infected bacteria and for the characterisation of the dynamics of infection, comprehensive and detailed studies are required to validate these observations and to fully understand their implications for characterizing bacteriophage-bacterial host interactions.

**Figure 5:**
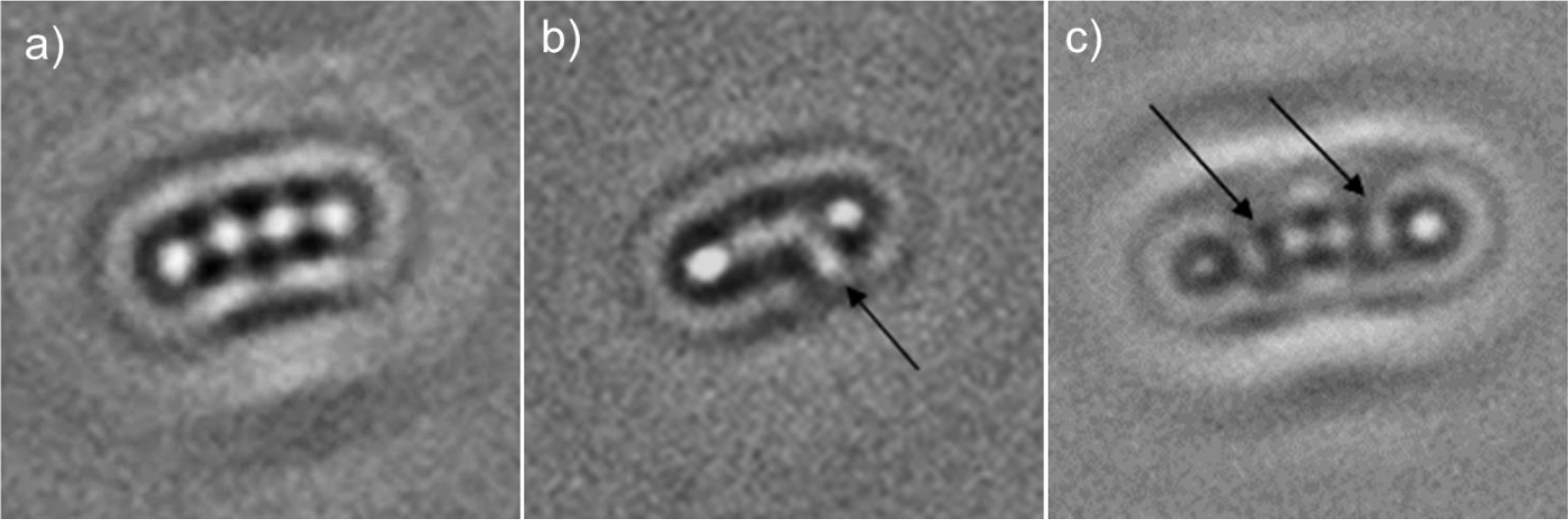
Details of E.Coli bacteria: a) not exposed versus b) and c) exposed to a population of EcoLiv25 phages in solution. In figure the arrow point at the supposed presence of a phage attached to the bacterium external membrane while in figure c) the arrows nt at the compromised sections of the bacterium external membrane as a result of phages infection.

## 5. Conclusion

In this study, we introduce a novel, label-free visualization technique for bacteriophages utilizing an optical setup based on caustics. Our results confirm the method’s effectiveness in visualizing a range of phages with morphologies and sizes, enabling the real-time observation of their dynamics without fluorescent labelling. By employing caustics-based microscopy, we overcome several limitations of conventional high-resolution microscopy techniques, such as the invasive sample preparation required by SEM, TEM, and AFM, and the challenges associated with fluorescence labelling, including photobleaching and potential behavioural alteration of phages in optical microscopy. This approach allows for the study of phages in vitro in an optical microscope, offering a less invasive avenue for exploring phage behaviour, dynamics and life cycles.

Caustics microscopy has been proven effective across phages with varying size and genomic characteristics. This is demonstrated by the optical signatures generated by genetically diverse phages of *Pseudomonas aeruginosa*, including a jumbo phage (phiKZ), as well as on phages able to target different species (EcoLiv25), highlighting the broad applicability of this microscopy technique across diverse phage collections. This method enables the precise monitoring and tracking of phage movement through specific environments, such as body fluids or various clinical settings. These capabilities are essential for translating phage research into practical applications that address real-world health challenges, particularly in the field of infection control and personalized medicine. Furthermore, caustics microscopy potentially facilitates real-time observation of interactions between phages and their bacterial hosts, providing novel insights into the infection process and the dynamic relationships between these microorganisms. This advance could support the development of phage-based applications in biotechnology and medicine, particularly in the design of new antimicrobial therapies and the use of phages in genetic engineering and as biocontrol agents.

## 7. Authors’ contributions

LB isolated the phage strains investigated in this study and prepared appropriate phage solutions for testing under the supervision of JF. SC supported the preparation of the phage solutions and acquired TEM images. FG conducted the microscopy experiments under the supervision of JMC and EAP. All authors contributed to the analysis and interpretation of the results. FG prepared the first draft of the manuscript, and all authors contributed to the production of the final version. All authors have given approval to the final version of the manuscript.

## 8. Acknowledgment

The authors would like to acknowledge Rosanna Wright (University of Manchester) for supplying phage phiKZ.

